# STAT2 signaling as double-edged sword restricting viral dissemination but driving severe pneumonia in SARS-CoV-2 infected hamsters

**DOI:** 10.1101/2020.04.23.056838

**Authors:** Robbert Boudewijns, Hendrik Jan Thibaut, Suzanne J. F. Kaptein, Rong Li, Valentijn Vergote, Laura Seldeslachts, Carolien De Keyzer, Lindsey Bervoets, Sapna Sharma, Johan Van Weyenbergh, Laurens Liesenborghs, Ji Ma, Sander Jansen, Dominique Van Looveren, Thomas Vercruysse, Dirk Jochmans, Xinyu Wang, Erik Martens, Kenny Roose, Dorien De Vlieger, Bert Schepens, Tina Van Buyten, Sofie Jacobs, Yanan Liu, Joan Martí-Carreras, Bert Vanmechelen, Tony Wawina-Bokalanga, Leen Delang, Joana Rocha-Pereira, Lotte Coelmont, Winston Chiu, Pieter Leyssen, Elisabeth Heylen, Dominique Schols, Lanjiao Wang, Lila Close, Jelle Matthijnssens, Marc Van Ranst, Veerle Compernolle, Georg Schramm, Koen Van Laere, Xavier Saelens, Nico Callewaert, Ghislain Opdenakker, Piet Maes, Birgit Weynand, Christopher Cawthorne, Greetje Vande Velde, Zhongde Wang, Johan Neyts, Kai Dallmeier

## Abstract

Since the emergence of SARS-CoV-2 causing COVID-19, the world is being shaken to its core with numerous hospitalizations and hundreds of thousands of deaths. In search for key targets of effective therapeutics, robust animal models mimicking COVID-19 in humans are urgently needed. Here, we show that productive SARS-CoV-2 infection in the lungs of mice is limited and restricted by early type I interferon responses. In contrast, we show that Syrian hamsters are highly permissive to SARS-CoV-2 and develop bronchopneumonia and a strong inflammatory response in the lungs with neutrophil infiltration and edema. Moreover, we identify an exuberant innate immune response as a key player in pathogenesis, in which STAT2 signaling plays a dual role, driving severe lung injury on the one hand, yet restricting systemic virus dissemination on the other. Finally, we assess SARS-CoV- 2-induced lung pathology in hamsters by micro-CT alike used in clinical practice. Our results reveal the importance of STAT2-dependent interferon responses in the pathogenesis and virus control during SARS-CoV-2 infection and may help rationalizing new strategies for the treatment of COVID-19 patients.

## Introduction and Results

SARS-CoV-2 belongs to the family of *Coronaviridae*, which contains a large group of viruses that are constantly circulating in animals and humans. Illness in humans caused by coronaviruses is mostly mild and manifested by respiratory or digestive problems as leading symptoms^1^. However, some coronaviruses, such as SARS-CoV-1, MERS-CoV and the recent SARS-CoV-2, have been responsible for serious outbreaks of severe and lethal respiratory disease^2,3^. Unlike the previous outbreaks with SARS-CoV-1 and MERS-CoV, the current SARS-CoV-2 outbreak has evolved to the largest global health threat to humanity in this century.

The unprecedented scale and rapidity of the current pandemic urges the development of efficient vaccines, antiviral and anti-inflammatory drugs. A key step in expediting this process is to have animal models that recapitulate and allow to understand viral pathogenesis, and that can in particular be used to identify new drug targets and preclinically assess preventive and therapeutic countermeasures.

Acute respiratory disease caused by SARS-CoV-1 and MERS infections is characterized by a dysregulated inflammatory response in which a delayed type I interferon (IFN) response promotes the accumulation of inflammatory monocyte-macrophages^4–6^. The severe lung disease in COVID-19 patients seems to result from a similar overshooting inflammatory response^7^. However, because even non-human primates do not fully replicate COVID-19, little information and no appropriate animal models are currently available to address this hypothesis^8^.

To address this knowledge gap, we compared the effect of SARS-CoV-2 infection in wild-type (WT) mice of different lineages (BALB/c and C57BL/6) and Syrian hamsters, as well as a panel of matched transgenic mouse and hamster strains with a knockout (KO) of key components of adaptive and innate immunity. We used an original patient isolate of SARS-CoV-2 (BetaCoV/Belgium/GHB-03021/2020) that was passaged on HuH7 and Vero E6 cells for these studies (Fig. S1 and Fig. S2A). For full characterization and to exclude possible contaminants, we performed deep sequencing on the inoculum that was used to infect the animals (Fig. S2A). No adventitious agents could be detected (data not shown). However, two in-frame deletions in the N-terminal domain and the furin-cleavage site of Spike (S) glycoprotein (9aa and 5aa, respectively) had occurred between cell culture passage P4 (mixed population of 85% WT genomes and 15% (9+5aa del) mutant genomes) and P6 (100% (9+5aa del) mutant genomes)^9–11^, likely as adaptation to growth in Vero E6 cells *in vitro* (Fig. S2B).

To first examine whether adaptive immunity contributed to the susceptibility to SARS-CoV-2 infection, we inoculated WT (immune-competent) and SCID mice (lacking functional T and B cells) from the same BALB/c background intranasally with a high 2 × 10^5^ TCID_50_ viral dose (P4 virus) (Fig. 1A). On day 3 p.i., a viral RNA peak in the lungs was observed (Fig. 1B and Fig. S3) with no obvious differences in viral loads (Fig. 1B) nor lung pathology (Fig. 1D and Fig. S4A and S4B) between WT and SCID mice. These data indicate that mice that lack the human ACE2 receptor^12^, can in principle be infected with SARS-CoV-2, although inefficiently and likely transiently, as also observed for SARS-CoV-1^4,13^. However, adaptive immunity did not markedly contribute to this low susceptibility.

**Figure 1.**
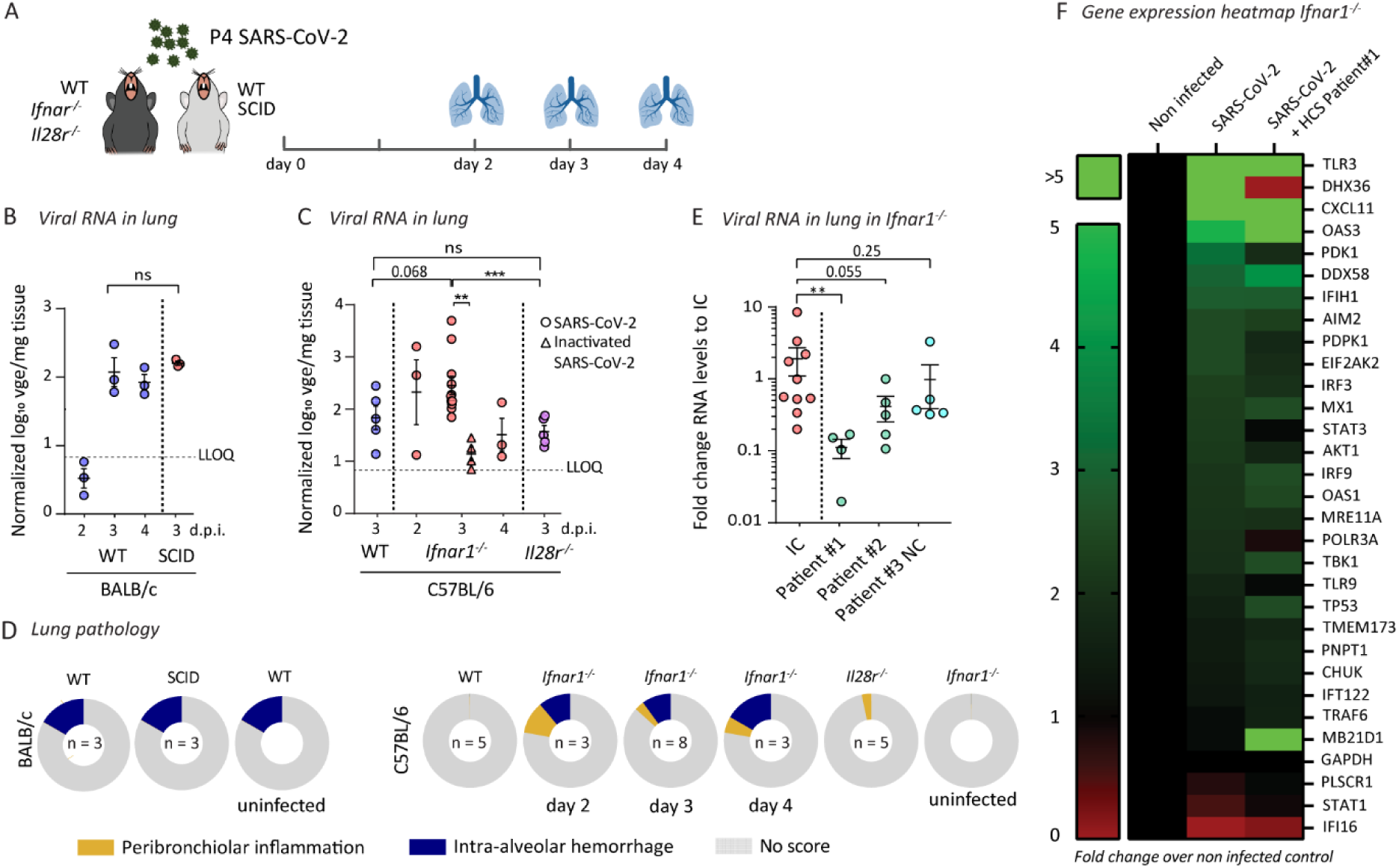
Type I interferon signaling restricts infection of the lungs of mice. (**A**) Schematic representation of SARS-CoV-2 inoculation schedule. Several wild-type (WT) and knock-out mouse strains were intranasally inoculated with 2 × 10^5^ TCID_50_ of passage 4 (P4) SARS-CoV- 2. On the indicated days post inoculation (d.p.i.), lungs were collected for determination of viral RNA levels and scored for lung damage. (**B-C**) Normalized viral RNA levels in the lungs of BALB/c WT (n=3) and SCID (n=3) mice and C57BL/6 WT (n=5), *Ifnar1*^*-/-*^ (n=3-8) and *Il28r*^*-/-*^ (n=5) mice. At the indicated time intervals p.i., viral RNA levels were determined by RT-qPCR, normalized against β-actin mRNA levels and transformed to estimate viral genome equivalents (vge) content per weight of the lungs (Figure S2). For heat-inactivation, SARS-CoV-2 was incubated for 30min at 56°C. Dotted line indicates lower limit of quantification (LLOQ). The data shown are means ± SEM. (**D**) Histopathological scoring of lungs for all different mouse strains. Mice were sacrificed on day 3 p.i. and lungs were stained with H&E and scored for signs of lung damage (inflammation and hemorrhage). Scores are calculated as percentage of the total maximal score. “No score” means not contributing to theoretical full cumulative score of 100%. Numbers (n) of animals analyzed per condition are given in the inner circle. (**E**) Viral RNA levels in *Ifnar1*^*-/-*^ mice after treatment with anti-SARS-CoV-2 serum or plasma. Mice were either left untreated (IC, infection control), or treated intraperitoneally one day before infection with convalescent serum (patient #1), convalescent plasma (patient #2) or with negative control plasma (patient #3 NC, negative control) and sacrificed on day 3 p.i. Viral RNA levels were determined in the lungs, normalized against β- actin and fold-changes were calculated using the 2^(-ΔΔCq)^ method compared to mean of IC. The data shown are means ± SEM. (**F**) Heatmap showing gene expression profiles of 30 selected marker genes in the lungs of uninfected and infected *Ifnar1*^*-/-*^ mice that were either left untreated or treated with convalescent serum from patient #1 (n=3 per group). Analysis performed on day 3 p.i. The scale represents fold change compared to non-infected animals. Statistical significance between groups was calculated by the nonparametric two-tailed Mann-Whitney U-test (ns = not significant, P > 0.05, * P < 0.05, ** P < 0.01, *** P < 0.001).

Interferons are the prototypic first-line innate immune defense against viral infections. To evaluate interferons, we compared viral RNA levels and lung pathology in WT C57BL/6 mice, and C57BL/6 mice with a genetic ablation of their type I (*Ifnar1*^*-/-*^) and III interferon (IFN) receptors (*Il28r*^-/-^) (Fig. 1A). *Ifnar1*^*-/-*^ mice showed an enhanced replication of SARS-CoV-2 in the lung on day 3 p.i. compared to both WT and *Il28r*^*-/-*^ mice (Fig. 1C). Similar to BALB/c mice, overall viral loads were low. *Ifnar1*^*-/-*^ mice that were treated prior to infection with human convalescent SARS-CoV-2 patient serum or plasma that contains spike-specific antibodies (Fig. 1E, Fig. S4) had a 3-10-fold reduction in viral loads depending on the patient donor. This provides further evidence for active, although inefficient virus replication in *Ifnar1*^*-/-*^ mice.

WT and knockout (*Ifnar1*^*-/-*^, *Il28r*^*-/-*^) mouse strains, all on C57BL/6 background, presented consistently with only a mild lung pathology. However, *Ifnar1*^*-/-*^ mice showed increased levels of intra-alveolar hemorrhage, sometimes accompanied by some peribronchiolar inflammation (Fig. 1D and Fig. S5A and S5B). Passive transfer of HCS did not result in an obvious improvement in histopathological scores (Fig. S5C), in line with other studies about partial protection from SARS-CoV-2 infection^14,15^ and virus-induced inflammatory responses. Further evidence for true infection and hence viral replication is provided by transcriptomic analysis (Sharma, S. *et al.*, in press^16^) of infected lung tissues (Fig. 1F and Fig. S6), revealing (i) an upregulation of classical antiviral effector molecules^17^ (enrichment p<0.001) such as *cGAS, Mx1, IFIH1/MDA-5, IRF3, OAS1, OAS3* and *PKR/EIF2AK2* (Fig. S6, Cluster 1) and (ii) downregulation of upstream regulators *STAT1, STAT3* and *STING/TMEM173* (Fig. S6, Cluster 3) in agreement with a possible role for apoptosis^18–20^. Likewise, HCS treatment modulated, at least to some extent, the observed gene expression patterns (Fig. 1F and Fig. S6, Cluster 2) as shown by decreasing *Akt1* (p=0.034) or increasing *DDX58* (*RIG-I*, p=0.028) and *cGAS* (*MB21D1*, p=0.094) mRNA levels. In summary, our data are in line with restriction of SARS-CoV-2 infection by the interferon system in mice, and also suggest limited inflammatory responses in the lungs of mice, in contrast to COVID-19 in humans^21^. Taken together, mice were considered as a poor model to study COVID-19 pathogenesis, or to assess the efficacy of vaccines and treatments.

In contrast, Syrian hamsters have been reported to be highly susceptible to SARS-CoV-1^22^ and SARS-CoV-2^23^ and might thus provide a small animal model to study SARS-CoV-induced pathogenicity and the involvement of the immune response in aggravating lung disease. In contrast to mice, intranasal inoculation of SARS-CoV-2 in WT hamsters resulted in high viral RNA loads (Fig. 2B, Fig. S7) a proxy used for the quantification of viral loads (see Fig. S9C), and in actual infectious titers (Fig. 2C) in the lungs, i.e. roughly 4 Log_10_ higher than in *Ifnar*^*-/-*^ mice (Fig. 2C). Also, a marked lung pathology [median cumulative score (MCS) 9 out of maximal score of 18; IQR=8.5-10.5 (P4 virus)] characterized by a multifocal necrotizing bronchiolitis, massive leukocyte infiltration and edema was observed in infected hamsters but not in mice (Fig. 2D and Fig. S8A-C). This resembles histopathological findings in humans suffering from severe bronchopneumonia^24^.

**Figure 2.**
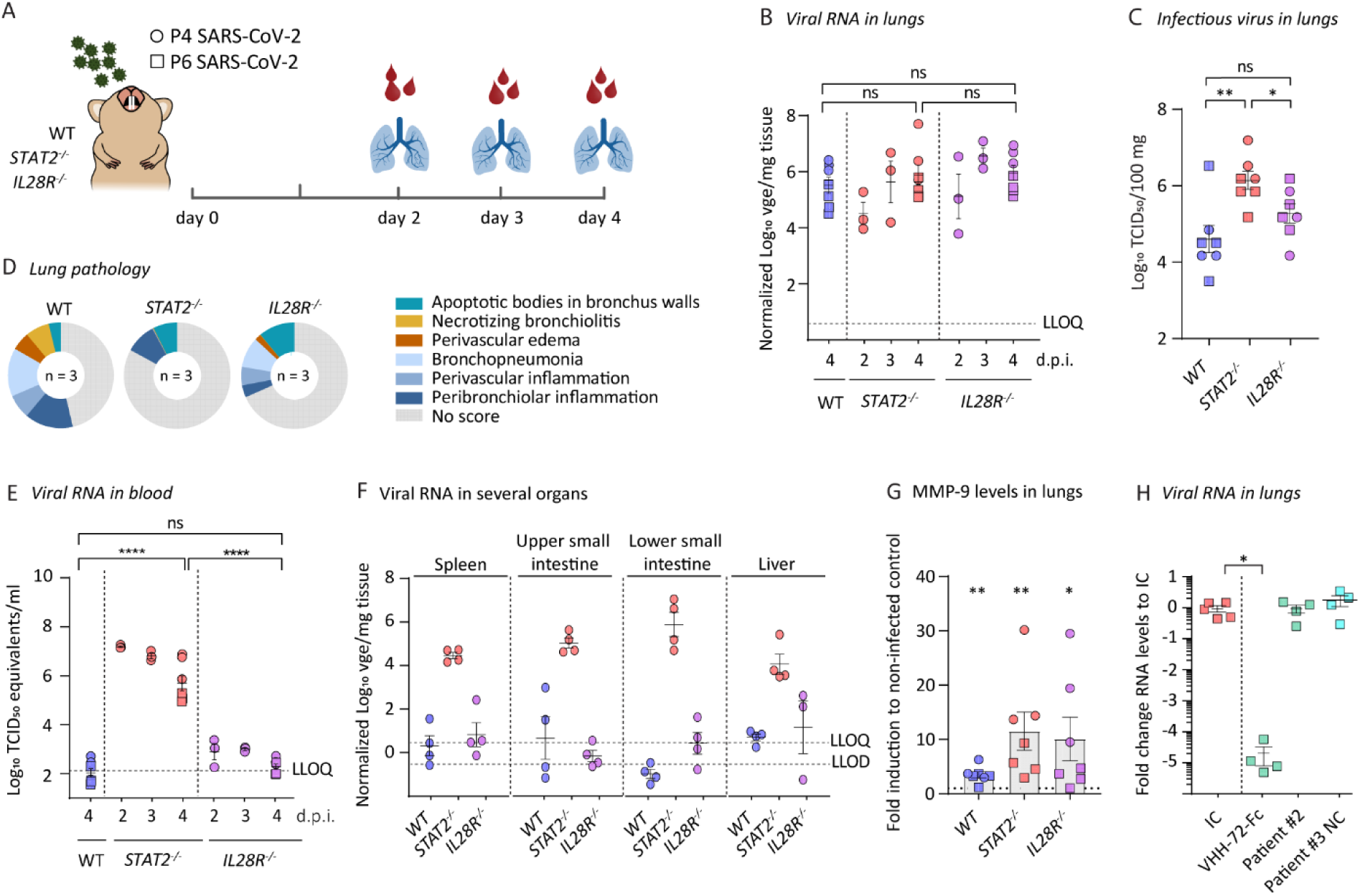
Exuberant innate response by *STAT2* drives SARS-CoV-2-induced lung pathology in hamsters. **(A)** Schematic representation of SARS-CoV-2 inoculation schedule. WT, *STAT2*^*-/-*^ and *IL28R-a*^*-/-*^ hamster strains were intranasally inoculated with 2 × 10^5^ TCID_50_ of passage 4 or 2 × 10^6^ of passage 6 SARS-CoV-2. Outcomes derived from inoculation with passage 4 or passage 6 SARS-CoV-2 is designated by circles (P4) or squares (P6). On the indicated days post inoculation (d.p.i.), organs and blood were collected to determine viral RNA levels, infectious viral load and score for lung damage. Viral loads in the indicated organs were quantified by RT-qPCR **(B, E and F)** or virus titration **(C). (B**,**F)** Viral RNA levels in the indicated organs were normalized against β-actin mRNA levels and transformed to estimate viral genome equivalents (vge) content per weight of the lungs (Figure S5)**. (C)** Infectious viral loads in the lung are expressed as the number of infectious virus particles per 100 mg of lung tissue. **(E)** Viral RNA levels in the blood were calculated from a standard of infectious virus and expressed as TCID_50_ equivalents per ml blood. Dotted lines indicate lower limit of quantification (LLOQ) or lower limit of detection (LLOD) **(D)** Histopathological scoring of lungs. Hamsters were sacrificed on day 4 p.i. with passage 4 SARS-CoV-2 and lungs were stained with H&E and scored for signs of lung damage (apoptotic bodies, necrotizing bronchiolitis, edema, pneumonia and inflammation). Scores are calculated as percentage of the total maximal score. **(G)** Levels of matrix metalloproteinase (MMP)-9 levels in lung homogenates of SARS-CoV-2 infected hamsters, relative to non-infected controls of the same strain. Statistical significance was calculated between infected and non-infected animals within each group. Values for infected animals (n=7 each) compiled from two independent experiments using either P4 (n=3, circles) and P6 (n=4, squares) SARS-CoV-2. **(H)** Viral RNA levels in hamsters after treatment with convalescent SARS-CoV-2 plasma or with a previously described antibody. Hamsters were either left untreated (IC, infection control, n=5) or treated with a single-domain antibody (VHH-72-Fc, n=4), convalescent plasma (patient #2, n=4) or negative control plasma (patient #3 NC, negative control, n=4) and sacrificed on day 4 p.i. Viral RNA levels were determined in the lungs, normalized against β-actin and fold-changes were calculated using the 2^(-ΔΔCq)^ method compared to the mean of IC. The data shown are means ± SEM. Statistical significance between groups was calculated by the nonparametric two-tailed Mann-Whitney U-test (ns P > 0.05, * P < 0.05, ** P < 0.01, **** P < 0.0001).

In order to investigate the roles of type I and III IFN in the pathogenesis of SARS-CoV-2 infection, we compared virus replication levels and lung pathology in WT hamsters and hamsters with ablated *Signal Transducer and Activator of Transcription 2* (*STAT2*^*-/-*^ lacking type I and III IFN signaling)^25,26^ and IL28R expression (*IL28R-a*^*-/-*^ lacking IFN type III signaling) (Fig. 2A). Of note, these receptor knockouts did not affect ACE2 expression in hamster lungs (Fig. S9A), while interferon-stimulated genes (ISG)^27^ such as MX-2 (strongly induced by IFNα/STAT2 signaling) and IP-10 (induced by both type I and type II IFNs) showed a differential expression pattern when comparing the different genotypes, triggered by SARS-CoV-2 infection (Fig. S9B). Importantly, lower baseline expression of MX-2 and IP-10 and failure to respond to SARS-CoV-2 infection by MX-2 upregulation in *STAT2*^*-/-*^ hamsters confirmed the functional knockout. As expected, *IL28R-a*^*-/-*^ hamsters showed an intermediate phenotype between that of WT and *STAT2*^-/-^ concerning their antiviral response. For many respiratory viruses, including SARS-CoV-1, type I and III interferon signaling has been described to play an important role in restricting infection^28^. No marked differences were observed in viral RNA levels in the lung of WT, *STAT2*^*-/-*^ or *IL28R-a*^*-/-*^ hamsters (Fig. 2B). However, *STAT2*^*-/-*^ hamsters had higher titers of infectious virus in the lung (Fig. 2C), high titer viremia (measured by RT-qPCR and virus titration)^29^ (Fig. 2E and Fig. S9C) and high levels of viral RNA in the spleen, liver and upper and lower gastrointestinal tract^30^ (Fig. 2F) in comparison with WT and *IL28R-a*^*-/-*^ hamsters. Interestingly, only for the P4 virus and not the P6 virus, infectious virus could be observed in the blood. Together, these data suggest STAT2 is critical for restricting SARS-CoV-2 systemic spread and suppressing viral replication outside of the lung compartment. A similar effect was also observed in human Calu-3 cells where pharmacological inhibition of STAT signaling by Ruxolitinib could rescue virus replication from the inhibitory activity of type I IFN (Fig. S9D), further underscoring the role of STAT2 in controlling virus replication. Inversely, the observed lung pathology was much attenuated in *STAT2*^*-/-*^ hamsters [MCS=3; IQR=1.5-3 (P4 virus)] with a limited infiltration of polymorphonuclear leukocytes correlated with the detection of few apoptotic bodies in the bronchus walls (Fig. 2D and Fig. S8C). On the contrary, *IL28R-a*^*-/-*^ hamsters showed clear signs of bronchopneumonia and peribronchiolar inflammation, yet of an intermediate score [MCS=7; IQR=6.5-7 (P4 virus)] (Fig. 2D and Fig. S8C). Matrix metalloprotease (MMP)-9 levels, which may serve as a sensitive marker for the infiltration and activation of neutrophils in inflamed tissues^31,32^, were markedly elevated in the lungs of all infected hamsters (Fig. 2G). However, higher MMP-9 levels were found in *STAT2*^*-/-*^ animals, thereby inversely correlating with the histological findings (Fig. 2D). In addition, biomarkers elevated in critically ill COVID-19 patients^2,7,33^ such as the cytokines IL-6, IL-10 and IFN-γ were not found to be elevated in the serum of infected hamsters (Fig. S10B), although mRNA levels of IP-10 (CXL-10) were upregulated in the lungs of SARS-CoV-2 infected hamsters as reported for other cytokine/chemokines downstream of IFN-γ ^23^ (Fig. S9B). Nonetheless, infected *STAT2*^*-/-*^ and *IL28R-a*^*-/-*^ had clearly increased levels of IL-6 and IL-10 in their lungs (Fig. S10A). Such an inverse correlation between biomarkers and pathology in WT versus *STAT2*^*-/-*^ hamsters is in line with findings in mouse models of SARS-CoV-1 infection in which pathology correlated with the induction and dysregulation of alternatively activated “wound-healing” monocytes/macrophages^4,6^. To assess the utility of the hamsters for testing the effect of therapeutic interventions on SARS-CoV-2 replication, WT hamsters were treated with human convalescent plasma or a neutralizing SARS-CoV-1 and SARS-CoV-2- specific single-domain antibody Fc fusion construct (VHH-72-Fc)^34^ prior to infection (Fig. 2H). Unlike a single dose of convalescent plasma, which did not significantly reduce viral load in the lungs, pre-treatment with VHH-72-Fc reduced viral loads in the lung ∼10^5^-fold compared to untreated control animals, validating hamsters as preclinical model for testing anti-SARS-CoV-2 therapies.

The lack of readily accessible serum markers or the absence of overt disease symptoms in hamsters prompted us to establish a non-invasive means to score for lung infection and SARS-CoV-2 induced lung disease by computed tomography (CT) as used in standard patient care to aid COVID-19 diagnosis with high sensitivity and monitor progression/recovery^7,33,35,36^. Similar as in humans^37^, semi-quantitative lung pathology scores were obtained from high-resolution chest micro-CT scans of free-breathing animals^38^. Pulmonary consolidations were present in SARS-CoV-2 infected WT and *IL28R- a*^*-/-*^ hamsters, but not in *STAT2*^*-/-*^ hamsters (Fig. 3A-B and Fig. S11A). In the upper airways, no differences were observed between WT and *STAT2*^*-/-*^ hamsters, whereas *IL28R-a*^*-/-*^ hamsters presented with an obvious dilation of bronchi (Fig. 3A and 3C). Further quantitative analysis^39^ revealed an increase of the non-aerated lung volume in SARS-CoV-2-infected WT and *IL28R-a*^*-/-*^ hamsters, yet again not in *STAT2*^*-/-*^ hamsters (Fig. 3D, Fig. S11B). Hence apart from lung consolidations and airway dilation as the main observed pathology, marked differences in other micro-CT-derived markers of specific lung pathology, such as hyperinflation, emphysema, or atelectasis^39,40^ were not be observed, except in one animal that presented with hyperinflation (Fig. S11C and Fig. S12A-C). A matched comparison of the micro-CT-derived lung scores and viral loads in the lungs showed that, in line with our previous results, type I IFN responses downstream of STAT2 signaling drive lung pathology, yet has minor impact on viral replication in the lungs (Fig. 3E). Together, these data fully support micro-CT as a convenient adjunct to histological scoring (Fig. 2D and Fig. S11D) to visualize and quantify SARS-CoV-2-induced lung injury in the hamster model. Moreover, it may allow to monitor the impact of therapeutic measures non-invasively during disease progression.

**Figure 3.**
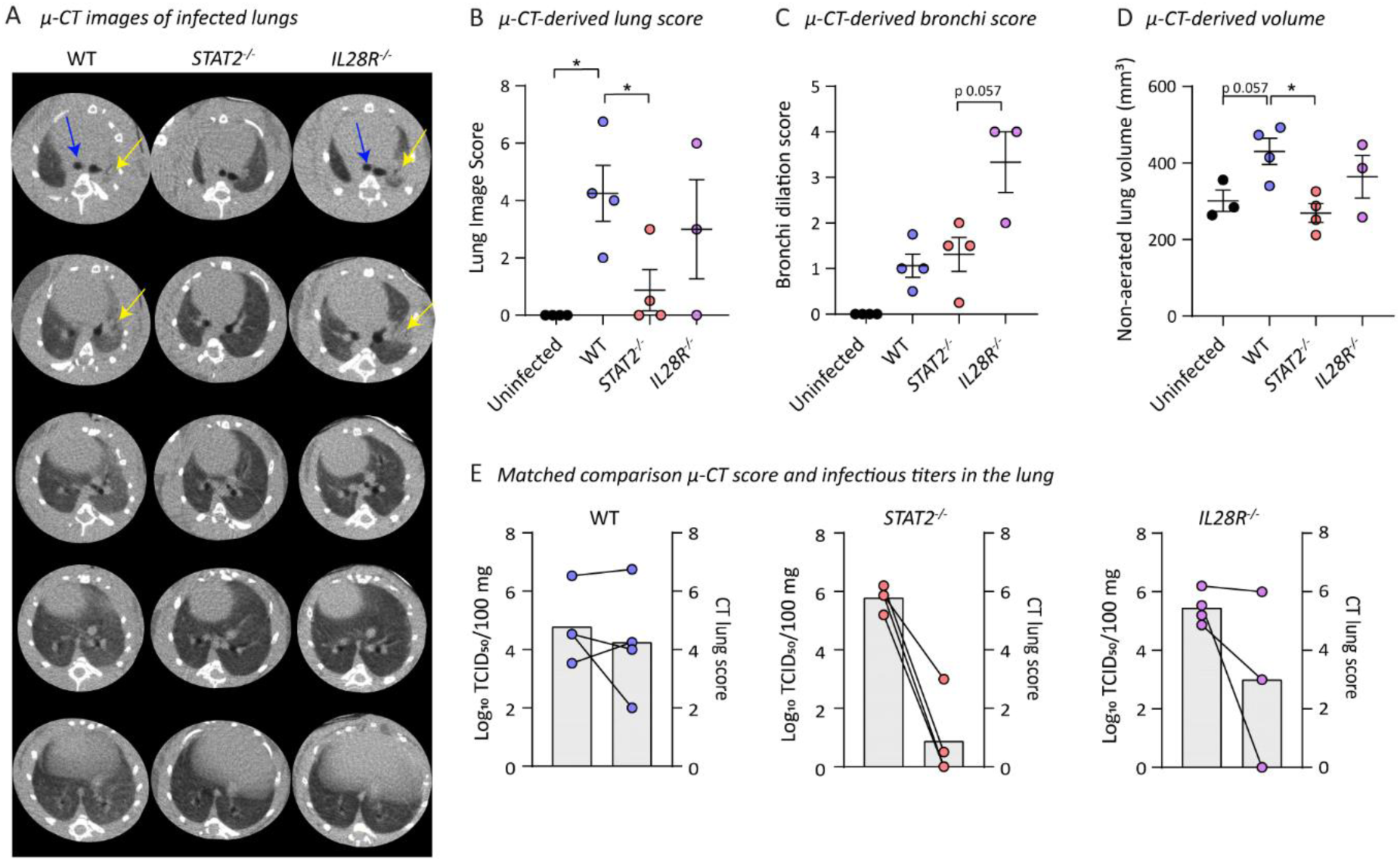
Micro-CT reveals severe lung injury in hamsters. **(A)** Representative transversal micro-CT images of infected (P6 SARS-CoV-2) WT (n=4), *STAT2*^*-/-*^ (n=4) and *IL28R-a*^*-/-*^ (n=3) hamster lungs at 4 d.p.i.. Arrows indicate examples of pulmonary infiltrates seen as consolidation of lung parenchyma (yellow) or dilatation of upper airways (blue). Five transverse cross sections at different positions in the lung were selected for each animal and scored to quantify lung consolidations **(B)** or dilatation of the bronchi **(C). (D)** Quantification of the micro-CT-derived non-aerated lung volume biomarker, reflecting the volume consolidations in the lungs. **(E)** Matched comparison between micro-CT-derived lung scores (left Y-axis) **(values from panel B of this figure)** and infectious viral load in the lung (right Y-axis) **(values from Fig. 2C)**. Lines indicate matched samples. The data shown are means ± SEM. Statistical significance between groups was calculated by the nonparametric two-tailed Mann Whitney U-test (ns P > 0.05, * P < 0.05).

## Discussion

The development of efficient therapeutic interventions against SARS-CoV-2 asks for relevant small animal models that mimic the different clinical manifestations of COVID-19 and that provide fundamental mechanistic insight into the underlying pathology/pathogenesis. Transgenic mice expressing *hACE2*, the *bona fide* receptor of SARS-CoV-1 and SARS-CoV-2^41^, have been suggested as COVID-19 model. We here demonstrate that WT mice are also susceptible to SARS-CoV-2 infection, yet resulting in very limited viral replication and inflammatory responses. Ablation of type I interferon signaling in *Ifnar*^*-/-*^ mice results in the same small incremental 10-fold increase in viral replication as was reported for SARS-CoV-2 in *hACE2* transgenic mice^41,42^. Most likely, neither mouse model will fully recapitulate pathogenesis of COVID-19, nor allow the study of clinical SARS-CoV-2 isolates. It remains of interest to investigate the added effect of both hACE2 knock-in and IFNAR knockout in mice.

By contrast, SARS-CoV-2 infection and associated pathology in hamsters seems to resemble what has been reported for SARS-CoV-1 in the same model. An early peak of active virus replication was noted in the lungs with viremia and extra-pulmonary spread. This was accompanied by a strong acute inflammatory response^22^ (as visualized by histopathology), the levels of which were correlated with those of MMP-9, a clinically relevant biomarker. Furthermore, micro-CT as established in this study may become a key instrument to non-invasively and quantitatively monitor SARS-CoV-2 lung disease. This will allow to conveniently monitor the effect of therapeutic strategies and test the preclinical efficacy of vaccine candidates. Unlike in humans, IL-6, IL-10 and IFN-γ levels were not elevated in SARS-CoV-2 infected hamsters. However, the immune response in COVID-19 patients is a dynamic process in which levels of these cytokine fluctuate over the different phases of disease often peaking late^7,43,44^. By contrast, hamsters show a very acute and short-lived infection^23^ that may not allow for the development of a similar cytokine storm that characterizes COVID-19 disease progression in humans, considering that in our studies animals were sacrificed as early as day 4 p.i. Nevertheless, the pathological findings in hamsters seem to resemble to a large extent what is observed in humans, including histopathology, infiltration of PMN cells and consolidation as scored by micro-CT.

By comparing data obtained from experiments using the P4 and P6 virus stocks, we observed a possible attenuated *in vivo* fitness of the P6 stock (lower MMP-9 levels and less infectious virus and viral RNA levels in blood) likely as a consequence of adaptation to enhanced growth in Vero cell cultures. This observation is also in line with what others have reported, namely that tissue culture-adaptive mutations similar to those we have observed (Fig. S2) may result in reduced fitness of SARS-CoV-2 in wild-type hamsters (no body weight loss and less severe pathology)^45^. These findings emphasize the possible consequences of using cell culture-adapted virus when studying SARS-CoV-2 infection in animal models.

By using unique knock-out hamster lines and human Calu-3 cells, we demonstrated that STAT2 plays a critical role in mediating antiviral responses and restricting systemic dissemination of SARS-CoV-2. This is in line with the effect of STAT1 in a mouse model of SARS-CoV-1 infection^46^. However and much in contrast to what is generally observed for viral infections in *Stat2*^*-/-*^ mice^47^ or *STAT2*^*-/-*^ hamsters^26,48,49^, the severe pathology induced by SARS-CoV-2 in WT hamsters is not observed in the absence of STAT2. Indeed, pneumonia as assessed by sensitive micro-CT was absent in *STAT2*^*-/-*^ hamsters. Considering the negative regulation of IL-6 and other mediators of inflammation by STAT2^47,50^, our hamster model may help to understand the immune pathogenesis of Acute Lung Injury (ALI) caused by highly pathogenic coronaviruses^22,28,51^ as well as other respiratory viruses^4^. Of note, a meta-analysis of infected human cells revealed a specific SARS-CoV-2-regulated transcriptional footprint for STAT2 that was much stronger than that of any other inflammatory transcription factor^52^.

The increase in replication of SARS-CoV-2 seen in *IL28R-a*^*-/-*^ hamsters, on one hand, combined with a tempered inflammatory response and lung injury as compared to WT hamsters, on the other hand, is in line with the role of type III IFN plays during respiratory virus infections, including SARS-CoV- 1^53^. This observation also suggests that in humans pegylated IFN-lambda^54,55^ (or similar modulators of innate immunity) may possibly be considered to protect medical staff and other frontline workers from SARS-CoV-2 infection or to dampen symptoms in critically ill patients^56^.

In conclusion, hamsters may be preferred above mice as infection model for the preclinical assessment of antiviral therapies, of convalescent serum transfer and of approaches that aim at tempering the COVID-19 immune pathogenesis in critically ill patients^21,57^. The latter may be achieved by repurposing anti-inflammatory drugs^58^ such as IL-6 receptor antagonists (e.g. Tocilizumab)^59^, or small molecule Jak/STAT inhibitors (e.g. Ruxolitinib or Tofacitinib). Educated by our finding that STAT2 signaling plays a dual role in also limiting viral dissemination, targeting the virus-induced cytokine response and overshooting of macrophage activation may need to be complemented by (directly acting) antivirals^60^.

## Methods

### Animals

Wild-type Syrian hamsters (*Mesocricetus auratus*) were purchased from Janvier Laboratories. All other mouse (C57BL/6, *Ifnar1*^*-/-*^, *Il28r*^*-/-*^, BALB/c and SCID) and hamster (*STAT2*^*-/-*^ and *IL28R-a*^*-/-*^) strains were bred in-house. Six- to eight-weeks-old female mice and wild-type hamsters were used throughout the study. Knock-out hamsters were used upon availability; seven- to twelve-week old female *STAT2*^*-/-*^ hamsters; five- to seven-week-old *IL28R-a*^*-/-*^ hamsters.

*Ifnar1*^*-/-*^ mouse breeding couples were a generous gift of Dr. Claude Libert, IRC/VIB, University of Ghent, Belgium. *Il28r*^*-/-*^ mice [C57B/6N-A<tm1Brd> Ifnlr1<tm1a(EUCOMM)Wtsi>/Wtsi, strain ID: EM:07988] were provided by the Wellcome Trust Sanger Institute Mouse Genetics Project (Sanger MGP)^61^.

*STAT2*^*-/-*^ and *IL28R-a*^*-/-*^ hamsters were generated by CRISPR/Cas-mediated gene targeting. To ablate *STAT2* (Gene ID: 101830537) expression, a 1-nt frameshift mutation was introduced in exon 4 resulting in multiple premature stop codons^25^; to ablate *IL28R* (*IFNLR1;* Gene ID: 101833778) expression, a 22-nucleotide deletion was introduced in exon 2 resulting in multiple premature stop codons in the original open reading frame.

Animals were housed individually (hamsters) or per 5 (mice) in individually ventilated isolator cages (IsoCage N Biocontainment System, Tecniplast) with access to food and water *ad libitum*, and cage enrichment (cotton and cardboard play tunnels for mice, wood block for hamsters). Housing conditions and experimental procedures were approved by the ethical committee of KU Leuven (license P015-2020), following institutional guidelines approved by the Federation of European Laboratory Animal Science Associations (FELASA). Animals were euthanized by 100µl (mice) or 500µl (hamsters) of intraperitoneally administered Dolethal (200mg/ml sodium pentobarbital, Vétoquinol SA). Animals were monitored daily for signs of disease (lethargy, heavy breathing or ruffled fur).

Prior to infection, the animals were anesthetized by intraperitoneal injection of a xylazine (16 mg/kg, XYL-M®, V.M.D.), ketamine (40 mg/kg, Nimatek, EuroVet) and atropine (0.2 mg/kg, Sterop) solution. Each animal was inoculated intranasally by gently adding 50µl droplets of virus stock containing 2 × 10^5^ TCID_50_ (P4 virus) or 2 × 10^6^ TCID_50_ (P6 virus) in both nostrils. Uninfected animals did not receive any virus or matrix. Due to time constraints by the pandemic outbreak and the urgency to develop a small animal model, we have used virus stock from two different passages (P4 and P6) to characterize our model. A full genotypic characterization of the viruses used is provided in Fig. S2, and phenotypic differences and virus titers are provided in the text and legends.

### Cells, virus and sera

Vero E6 (African green monkey kidney, kind gift from Peter Bredenbeek, LUMC, NL) and HuH7 (human hepatoma, JCRB0403) cells were maintained in minimal essential medium (Gibco) supplemented with 10% fetal bovine serum (Integro), 1% bicarbonate (Gibco), and 1% L-glutamine (Gibco). For maintenance of Calu-3 cells (human airway epithelium, kind gift from Lieve Naesens, KU Leuven, BE), the above medium was supplemented with 10mM HEPES (Gibco). All assays involving virus growth were performed using 2% (Vero E6 and HuH7) or 0.2% (Calu-3) fetal bovine serum instead of 10%.

SARS-CoV-2 strain BetaCov/Belgium/GHB-03021/2020 (EPI ISL 407976|2020-02-03) recovered from a nasopharyngeal swab taken from a RT-qPCR-confirmed asymptomatic patient returning from Wuhan, China beginning of February 2020^62^ was directly sequenced on a MinION platform (Oxford Nanopore) as described previously^63^. Phylogenetic analysis confirmed a close relation with the prototypic Wuhan-Hu-1 2019-nCoV (GenBank accession number MN908947.3) strain. Infectious virus was isolated by serial passaging on HuH7 and Vero E6 cells (see Fig. S1), with the addition of penicillin/streptomycin, gentamicin and amphotericin B. Virus used for animal experiments was from passages P4 and P6. Prior to inoculation of animals, virus stocks were confirmed to be free of mycoplasma (PlasmoTest, InvivoGen) and other adventitious agents by deep sequencing on a MiSeq platform (Illumina) following an established metagenomics pipeline^64,65^. The infectious content of virus stocks was determined by titration on Vero E6 cells by the Spearman-Kärber method. All virus-related work was conducted in the high-containment BSL3+ facilities of the KU Leuven Rega Institute (3CAPS) under licenses AMV 30112018 SBB 219 2018 0892 and AMV 23102017 SBB 219 2017 0589 according to institutional guidelines.

Human convalescent serum (HCS) was donated under informed consent (Patient #1). Human convalescent plasma (Patient #2) was obtained from Biobank Rode Kruis-Vlaanderen, registered under Belgian law as Biobank BB190034 (Fig. S4). Plasma donated by a healthy volunteer sampled prior to emergence of SARS-CoV-2 served as negative control (NC donor). Serum/plasma was administered i.p. 1 day prior to infection, in a volume of 200µl per mouse and 1000µl per hamster. Antibody VHH-72-Fc was administered i.p. at a dose of 20mg/kg 1 day prior to infection. VHH-72- Fc was expressed in ExpiCHO cells (ThermoFisher Scientific) and purified from the culture medium as described^34^. Briefly, after transfection with pcDNA3.3-VHH-72-Fc plasmid DNA, followed by incubation at 32°C and 5% CO_2_ for 6-7 days, the VHH-72-Fc protein in the cleared cell culture medium was captured on a 5 mL MabSelect SuRe column (GE Healthcare), eluted with a McIlvaine buffer pH 3, neutralized using a saturated Na_3_PO_4_ buffer, and buffer exchanged to storage buffer (25 mM L-Histidine, 125 mM NaCl). The antibody’s identity was verified by protein- and peptide-level mass spectrometry.

### RNA extraction and RT-qPCR

Animals were euthanized at different time-points post-infection, organs were removed and lungs were homogenized manually using a pestle and a 12-fold excess of cell culture medium (DMEM/2%FCS). RNA extraction was performed from homogenate of 4 mg of lung tissue with RNeasy Mini Kit (Qiagen), or 50µl of serum using the NucleoSpin kit (Macherey-Nagel), according to the manufacturer’s instructions. Other organs were collected in RNALater (Qiagen) and homogenized in a bead mill (Precellys) prior to extraction. Of 100µl eluate, 4µl was used as template in RT-qPCR reactions. RT-qPCR was performed on a LightCycler96 platform (Roche) using the iTaq Universal Probes One-Step RT-qPCR kit (BioRad) with primers and probes (Table S1) specific for SARS-CoV- 2, mouse β-actin (*Actb*) and hamster β-actin (*ACTB*), *ACE2, MX2* and *IP-10* (IDT). For each data point, qPCR reactions were carried out in duplicate. Standards of SARS-CoV-2 cDNA (IDT) and infectious virus were used to express the amount of RNA as normalized viral genome equivalent (vge) copies per mg tissue, or as TCID_50_ equivalents per mL serum, respectively. The mean of housekeeping gene β-actin was used for normalization. The relative fold change was calculated using the 2^-ΔΔCt^ method^66^.

### Quantification of SARS-CoV-2 infectious particles in lung tissues

After extensive transcardial perfusion with PBS, lungs were collected, extensively homogenized using manual disruption (Precellys24) in minimal essential medium (5% w/v) and centrifuged (12,000 rpm, 10min, 4°C) to pellet the cell debris. Infectious SARS-CoV-2 particles were quantified by means of endpoint titrations on confluent Vero E6 cell cultures. Viral titers were calculated by the Spearman-Kärber method and expressed as the 50% tissue culture infectious dose (TCID_50_) per 100mg tissue.

### Differential gene expression and bioinformatics analysis

To study differential gene expression, RNA was extracted from lung tissues using Trizol, subjected to cDNA synthesis (High Capacity cDNA Reverse Transcription Kit, Thermo Fisher Scientific), and qPCR using a custom Taqman qRT-PCR array (Thermo Fisher Scientific) of 30 genes known to be activated in response to virus infection^16^, as well as two housekeeping genes (Table S2). Data collected were analysed using the Quant Studio Design and Analysis (version 1.5.1) and Data Assist software (version 3.01, Thermo Fischer Scientific). Pathway, GO (Gene Ontology) and transcription factor target enrichment analysis was performed using GSEA (Gene Set Enrichment Analysis, Molecular Signatures Database (MSigDB), Broad Institute). Principal component analysis, correlation matrices, unsupervised hierarchical clustering (Eucledian distance) were performed using XLSTAT and visualized using MORPHEUS (https://software.broadinstitute.org/morpheus) as described previously^20^.

### Histology

For histological examination, the lungs were fixed overnight in 4% formaldehyde and embedded in paraffin. Tissue sections (4 µm) were stained with hematoxylin and eosin to visualize and score for lung damage.

### In vitro JAK/STAT inhibition assay

Calu-3 (human airway epithelial) cells were plated at 5×10^4^ cells/well in a 96-well plate and incubated overnight with 4µM of the JAK1/2 inhibitor Ruxolitinib^67^ (Toronto Research Chemicals, ON, Canada). Next day, cells were pre-treated for 4 hours with 10 IU/ml of Universal Type I IFN (PBL Assay Science, NJ, USA, cat no. 11200) before infection with SARS-CoV-2 (P6, 5×10^4^ TCID_50_ per well). Two hours post infection, cells were washed with PBS and incubated for an additional 48h with IFN and Ruxolitinib before collection of the supernatant for RNA extraction and quantification of virus yields by RT-qPCR.

### Tissue and serum biomarker analysis

Cytokine levels in lung homogenates and serum of hamsters were determined by ELISA for IFN-γ (EHA0005), IL-6 (EHA0008) and IL-10 (EHA0006) following the manufacturer’s instructions (Wuhan Fine Biotech Co., Ltd).

The levels of gelatinase B/metalloproteinase (MMP)-9 present in lung homogenates were analyzed using gelatin zymography^68^, essentially as described previously^69^. For quantification of zymolytic bands internal control samples were spiked into each sample. Equivalent hamster enzyme concentrations were calculated with the use of known amounts of recombinant human pro-MMP-9 and recombinant human pro-MMP-9ΔOGHem as standards^70^.

### Micro-computed tomography (CT) and image analysis

Hamsters were anaesthetized using isoflurane (Iso-Vet) (2-3% in oxygen) and installed in prone position into the X-cube micro-CT scanner (Molecubes) using a dedicated imaging bed. Respiration was monitored throughout. A scout view was acquired and the lung was selected for a non-gated, helical CT acquisition using the High-Resolution CT protocol, with the following parameters: 50 kVp, 960 exposures, 32 ms/projection, 350 µA tube current, rotation time 120 s. Data were reconstructed using a regularized statistical (iterative) image reconstruction algorithm using non-negative least squares^71^, using an isotropic 100 µm voxel size and scaled to Hounsfield Units (HUs) after calibration against a standard air/water phantom. The spatial resolution of the reconstruction was estimated at 200 µm by minimizing the mean squared error between the 3D reconstruction of the densest rod in a micro-CT multiple density rod phantom (Smart Scientific) summed in the axial direction and a digital phantom consisting of a 2D disk of 17.5 mm radius that was post-smoothed with Gaussian kernels using different full width half maxima (FWHM), after aligning the symmetry axis of the rod to the z-axis.

Visualization and quantification of reconstructed micro-CT data was performed with DataViewer and CTan software (Bruker micro-CT). As primary outcome parameter, a semi-quantitative scoring of micro-CT data was performed as previously described^38,39,72^ with minor modifications towards optimization for COVID-19 lung disease in hamsters. In brief, visual observations were scored (from 0 – 2 depending on severity, both for parenchymal and airway disease) on 5 different, predefined transversal tomographic sections throughout the entire lung image for both lung and airway disease by two independent observers (L.S. and G.V.V.) and averaged. Scores for the 5 sections were summed up to obtain a score from 0 to 10 reflecting severity of lung and airway abnormalities compared to scans of healthy, WT control hamsters. As secondary measures, image-derived biomarkers (non-aerated lung volume, aerated lung volume, total lung volume, the respective densities within these volumes and large airways volume) were quantified as in^38,72^ for a manually delineated VOI in the lung, avoiding the heart and main blood vessels. The threshold used to separate the airways and aerated (grey value 0-55) from non-aerated lung volume (grey value 56-255) was set manually on an 8-bit greyscale histogram and kept constant for all data sets.

### Statistical analysis

GraphPad Prism Version 8 (GraphPad Software, Inc.) was used for all statistical evaluations. The number of animals and independent experiments that were performed is indicated in the legends to figures. Statistical significance was determined using the non-parametric Mann Whitney U-test unless mentioned otherwise. Values were considered significantly different at P values of ≤0.05.

## Acknowledgments

We thank Kathleen Van den Eynde, Eef Allegaert, Sarah Cumps, Wilfried Versin, Caroline Collard, Elke Maas, Jasper Rymenants, Jasmine Paulissen, Nathalie Thys, Céline Sablon, Catherina Coun and Madina Rasulova for excellent technical assistance. We thank Johan Nuyts (Nuclear Medicine and Molecular Imaging, KU Leuven) for support with imaging file processing, Pieter Mollet (Molecubes, Ghent, Belgium) for outstanding technical support with the micro-CT installation, Jef Arnout and Annelies Sterckx (KU Leuven Faculty of Medicine, Biomedical Sciences Group Management) and Animalia and Biosafety Departments of KU Leuven for facilitating the studies. We also thank Dr. Claude Libert (IRC/VIB, University of Ghent) for providing the *Ifnar*^*-/-*^ mice and the Wellcome Trust Sanger Institute Mouse Genetics Project (Sanger MGP) and its funders for providing the *Il28r*^*-/-*^ mutant mouse line. We thank Berend Jan Bosch and Wentao Li (Utrecht University) for the Spike expression plasmid.

This project has received funding from the European Union’s Horizon 2020 research and innovation program under grant agreements No 101003627 (SCORE project) and No 733176 (RABYD-VAX consortium), funding from Bill and Melinda Gates Foundation under grant agreement INV-00636, and was supported by the Research Foundation Flanders (FWO) under the Excellence of Science (EOS) program (VirEOS project 30981113), the FWO Hercules Foundation (Caps-It infrastructure), the KU Leuven Rega Foundation, and the Stichting Antoine Faes. J.M. was supported by a grant from the China Scholarship Council (CSC). J.M.C. was supported by a doctoral grant from HONOURs Marie-Sklodowska-Curie training network (721367). B.V. is supported by a FWO SB grant for strategic basic research of the “Fonds Wetenschappelijk Onderzoek”/Research foundation Flanders [1S28617N]. This work is also supported by ‘Interne Fondsen KU Leuven / Internal Funds KU Leuven’ awarded to P.M. (project 3M170314). C.C. was supported by the FWO (FWO 1001719N). G.V.V. acknowledges grant support from KU Leuven Internal Funds (C24/17/061) and K.D. grant support from KU Leuven Internal Funds (C3/19/057 Lab of Excellence). G.O. and E.M. were supported by funding from KU Leuven (C16/17/010) and from FWO-Vlaanderen. D.D.V. was supported by a FWO sb fellowship and B.S. by FWO-EOS project VIREOS. COVID-19 research in the Saelens and Callewaert labs is supported by VIB, by a Ghent University GOA grant, and VIB, UGent and FWO emergency Covid-19 grants.

## Author Contributions

R.B., H.J.T., J.N. and K.D. designed experiments;

R.B., S.J.F.K, R.L., V.V., L.L., J.V.W., C.D.K., S.S., D.V.L., T.V., X.W., E.M., K.R., D.D.V., B.S., L.B., T.V.B., J.M., L.Cl., J.M.C., B.V., T.W.B., P.M. and W.C. carried out experiments;

R.B., H.J.T., L.S., S.J., J.V.W., D.J., E.M., B.W., P.M., C.C., G.V.V., Z.W. and K.D. analyzed data;

L.D., J.R.P., J.M., G.S., K.V.L., and G.O. provided advice on the interpretation of data;

R.B., H.J.T., and K.D wrote the original draft with input from co-authors;

R.B., H.J.T., L.S., J.V.W., C.C., G.V.V., J.N. and K.D. wrote the final draft;

R.L., Y.L., Z.W. developed the *STAT2*^*-/-*^ and *IL28R-a*^*-/-*^ hamster strains;

K.R., D.D.V., B.S., P.M., R.L., Y.L., Z.W., X.S., N.C., V.C., M.V.R. and G.O. provided essential reagents;

E.H., D.S., and P.L. provided and facilitated access to essential infrastructure;

H.J.T., J.N. and K.D. supervised the study;

K.D., L.Co., P.L. and J.N. acquired funding;

All authors approved the final manuscript.

## Data Availability Statement

All data supporting the findings in this study are available from the corresponding author upon request.

