## Supplementary Material for "STAT2 signaling as double-edged sword restricting viral dissemination but driving severe pneumonia in SARS-CoV-2 infected hamsters"

### Phylogenetic tree isolate

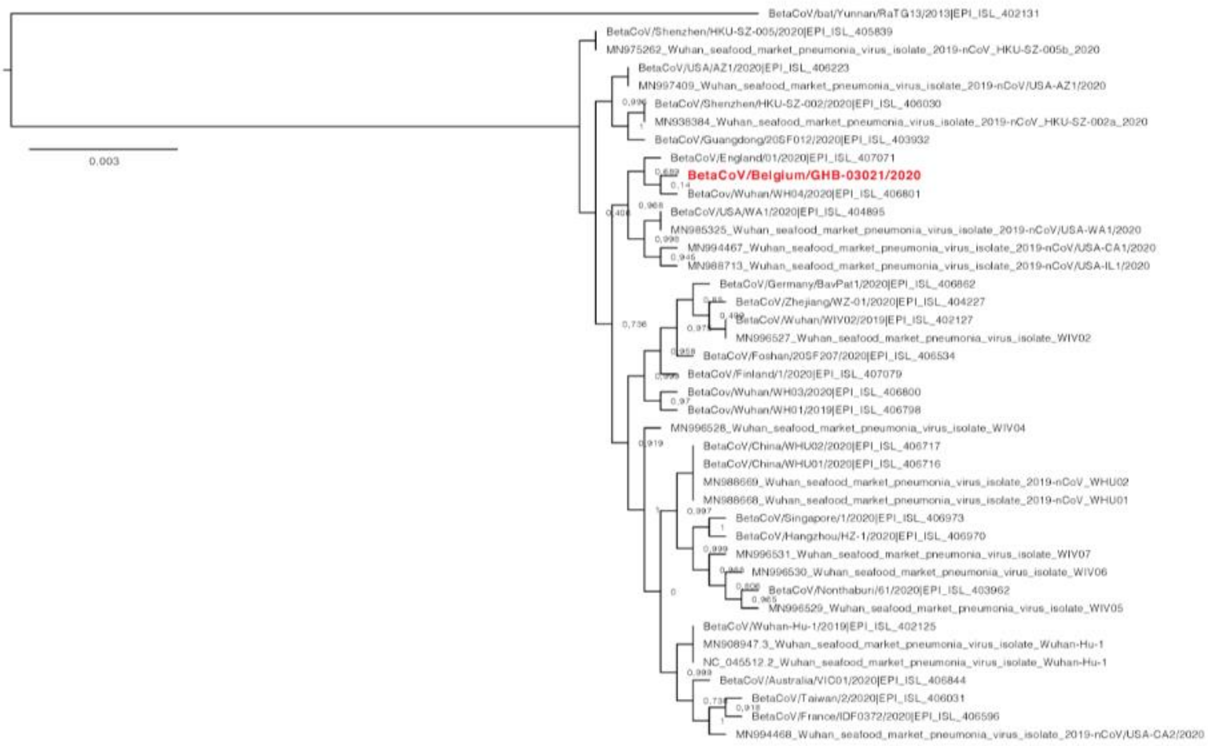


**Suppl Fig. S1. SARS-CoV-2 phylogeny.** Phylogenetic tree of recent clinical SARS-CoV-2 isolates, with the SARS-CoV-2 strain used in the current study indicated in red. Phylogenetic analysis confirmed a close evolutionary relationship of the isolate BetaCoV/Belgium/GHB-03021/2020 (EPI ISL 407976|2020-02-03) with the prototypic Wuhan-Hu-1 2019-nCoV strain (GenBank accession number MN908947.3).

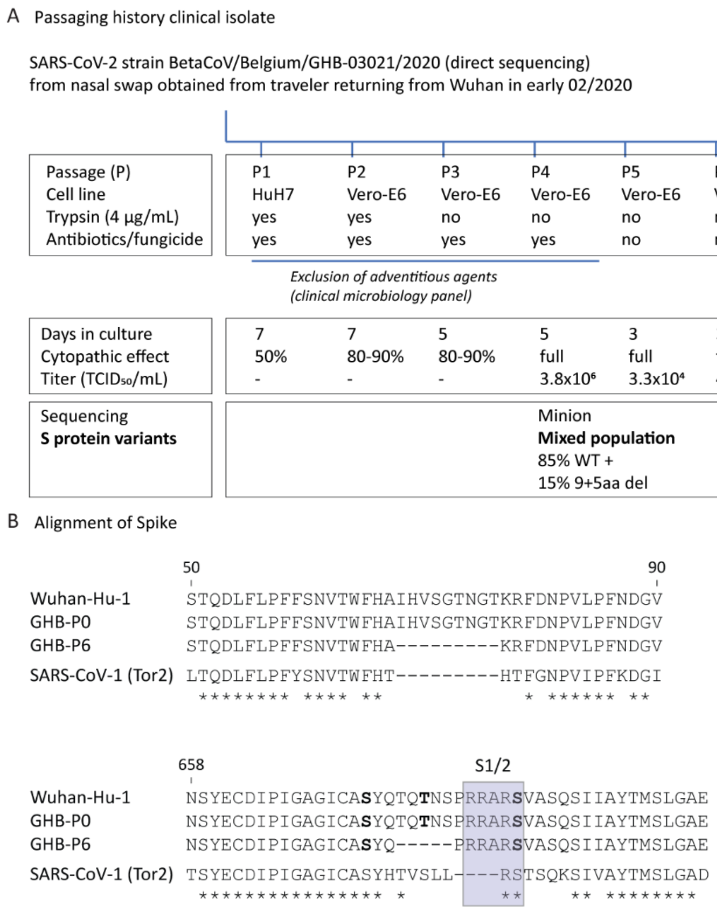


**Suppl Fig. 2 SARS-CoV-2 stock generation**. **(A)** Passaging (P) history of the SARS-CoV-2 isolate used indicating the origin and key characteristics of the virus stocks P4 and P6. Two in-frame deletions in the N-terminal domain and the furin-cleavage site of Spike (S) glycoprotein (9aa and 5aa, respectively) were observed in P4 (mixed population of 85% WT genomes and 15% (9+5aa del) mutant genomes). P6 only contains the (9+5aa del) mutant genomes. As a consequence, P4 is an intermediate between the original P0 WT virus and the P6 virus. **(B)** Alignment of parts of the Spike protein of SARS-CoV-2 isolates P0 and P6 with those of the prototypic Wuhan-Hu-1 strain[1,2], and SARS-CoV-1 (Tor2 strain)[3]. Upper panel – deletion discovered in the N-terminal domain (NTD); lower panel – deletion at the polybasic furin recognition site (shaded box)[4,5]; * – residue conserved between SARS-CoV-1 and SARS-CoV-2; **BOLD** – predicted O-glycosylation acceptor[6]; S1/S2 – junction between the membrane-distal S1 and membrane-proximal S2 subunits.

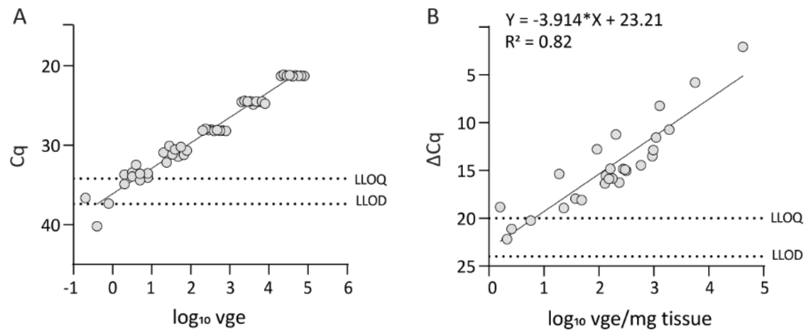


**Suppl Fig. S3. Correlation curves for quantification of viral loads in mice. (A)** Correlation between cycle of quantification (Cq) and $\log_{10}$ viral genome equivalents (vge) obtained from a cDNA plasmid standard. **(B)** Correlation between delta Cq and $\log_{10}$ vge/mg lung tissue in mice. Delta Cq values are calculated by subtracting Cq values of β-actin from Cq values of SARS-CoV-2. To express viral loads in the lungs of mice, this correlation was used to transform delta Cq values into *Normalized log10 vge/mg tissue*. Lowest limit of quantification (LLOQ) was defined as lowest detectable value in the linear range of the calibration curve. Lowest limit of detection (LLOD) was defined as lowest detectable value below the linear range of the calibration curve.

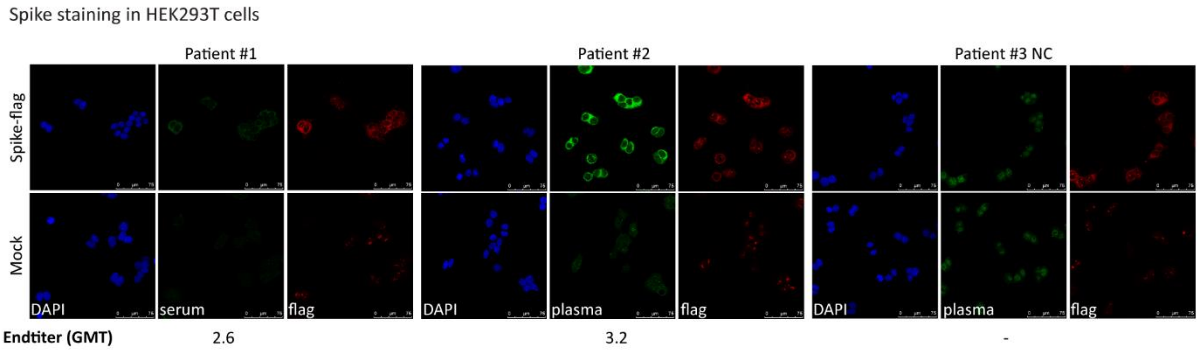


**Suppl Fig. S4. Spike-specific indirect immune fluorescence staining by convalescent donor blood.** HEK293T cells were transfected with a Spike-flag expression plasmid or mock transfected. One day after transfection, cells were fixed, permeabilized and stained with serum (patient #1) or plasma (patient #2 and #3 negative control, NC) from several human donors using specific fluorescence-labelled anti-human (green) and anti-FLAG (red) secondary antibodies. Fluorescence signal intensities were read out by high content imaging on a CX5 system (Thermo Scientific). Patients #1 and #2 have previously been diagnosed with SARS-CoV-2 whereas patient #3 is the negative control. Both patients #1 and #2 show Spike-specific staining that is not visible in the mock control and that correlates with FLAG staining. End titers ($log_{10}$ geometric mean titer, GMT) was defined as the last dilution in which a specific staining in the Spike-flag transfected cells was observed compared to the mock-transfected cells.

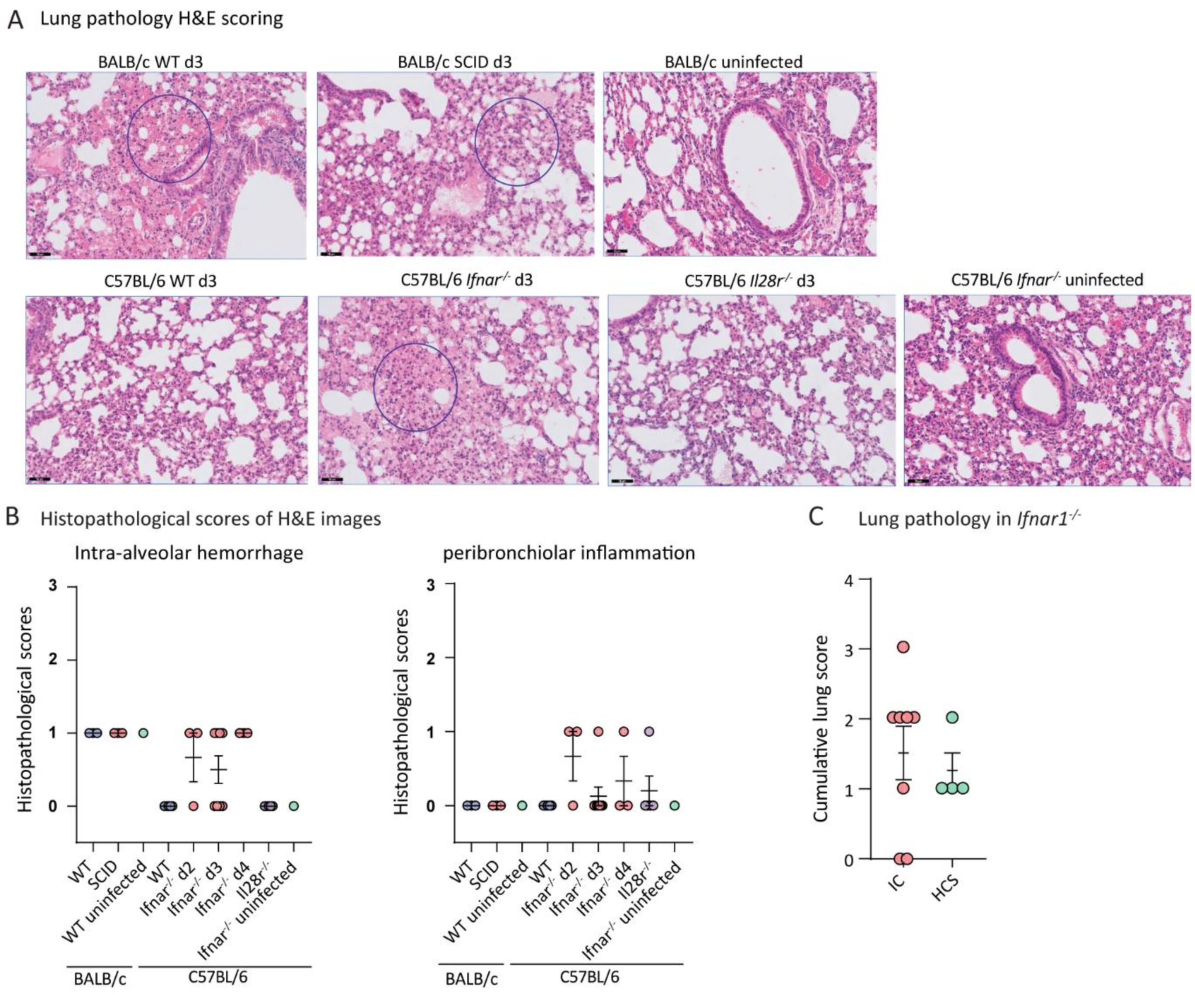


**Suppl Fig. S5. Lung pathology in mice. (A)** Representative H&E images of the lungs of uninfected or SARS-CoV-2 infected WT (n=3) and SCID (n=3) BALB/c and WT (n=5), *Ifnar1*$^{-/-}$ (n=3-8) and *Il28r*$^{-/-}$ (n=5) C57BL/6 mice. Circles indicate presence of intra-alveolar hemorrhage. **(B)** Histopathological scoring on a scale of 0-3 of intra-alveolar hemorrhage and peribronchiolar inflammation (WT – blue circles; SCID or Ifnar$^{-/-}$ – red; Il28r$^{-/-}$ – purple; uninfected controls – green) **(C)** Histopathological scoring of the lungs of *Ifnar1*$^{-/-}$ mice after treatment with anti-SARS-CoV-2 sera. Mice were either left untreated (IC, infection control) (n=8) or treated with HCS (human convalescent serum) (n=4) and sacrificed on day 3 p.i. Cumulative clinical scores of intra-alveolar hemorrhage and peribronchiolar inflammation are indicated. The data shown are means ± SEM. Statistical significance between groups was calculated by the nonparametric two-tailed Mann Whitney U-test (ns P > 0.05).

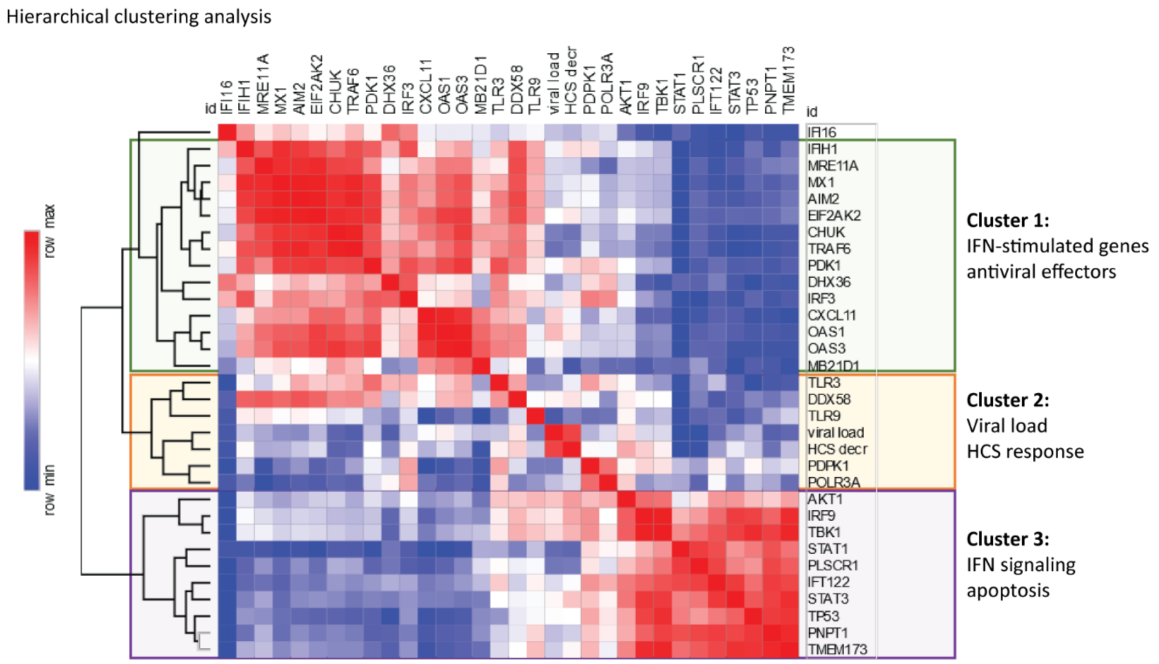


**Suppl Fig. S6. Hierarchical clustering analysis of transcriptomic data from uninfected and SARS-CoV-2-infected *Ifnar1*$^{-/-}$ mice**. Cluster dendrogram (Eucledian distance) and heatmap of Spearman correlation coefficients between molecular and virological data from uninfected and SARS-CoV-2-infected *Ifnar1*$^{-/-}$ mice, in the absence or presence of convalescent serum of patient #1 (HCS, human convalescent serum) (n=3 each, analysis on samples of day 3 p.i.), based on a transcriptomic analysis of 30 selected marker genes[7] (Table S2). HCS treatment modulates the gene expression pattern in SARS-CoV-2-infected lungs (Fig. 1F and Fig. S5) as shown by decreasing *Akt1* (p=0.034) and *IRF9* (p=0.076) or increasing *DDX58* (*RIG-I*, p=0.028) and *cGAS* (*MB21D1*, p=0.094) mRNA levels. Further hierarchical cluster analysis reveals three main clusters as indicated. A small intermediate cluster (Cluster 2) containing nucleic acid sensors *TLR3/TLR9/DDX58/POLR3A* correlates with viral load and HCS response. Clusters 1 and 3 are negatively correlated to each other and independent of viral load and HCS. Cluster 1 is enriched (p<0.001) for antiviral effector molecules such as *cGAS*, *Mx1*, *IFIH1/MDA-5*, *IRF3*, *OAS1*, *OAS3* and *PKR/EIF2AK2*[8]. Cluster 3 comprises upstream regulators *STAT1*, *STAT3* and *STING/TMEM173* but also *PLSCR1* (scramblase) and *p53*, linking IFN signaling to apoptosis (enrichment p<0.001)[9]. In contrast, *IFI16* decreases upon SARS-CoV-2 infection and clusters independently, suggesting absence of inflammatory pyroptosis, in line with a fast control of viral replication and limited tissue damage as observed in mice.

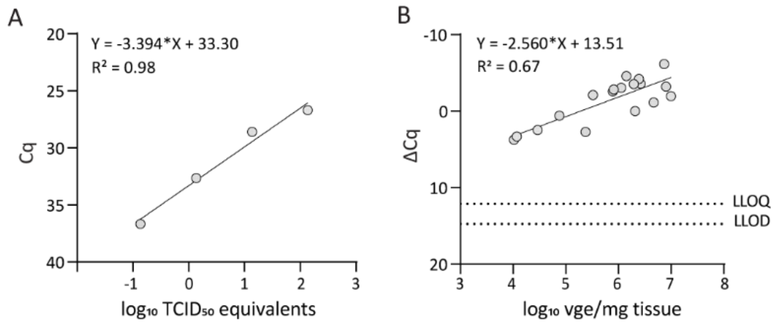


**Suppl Fig. S7. Correlation curves for quantification of viral loads in hamsters. (A)** Correlation between cycle of quantification (Cq) and $\log_{10}$ $TCID_{50}$ equivalents obtained from an infectious virus standard. **(B)** Correlation between delta Cq and log10 vge/mg tissue in hamsters. Delta Cq values are calculated by subtracting Cq values of β-actin from Cq values of SARS-CoV-2. To express viral loads in the different tissue of hamsters, this correlation was used to transform delta Cq values into *Normalized log10 vge/mg tissue*. Lowest limit of quantification (LLOQ) was defined as lowest detectable value in the linear range of the calibration curve. Lowest limit of detection (LLOD) was defined as lowest detectable value below the linear range of the calibration curve.

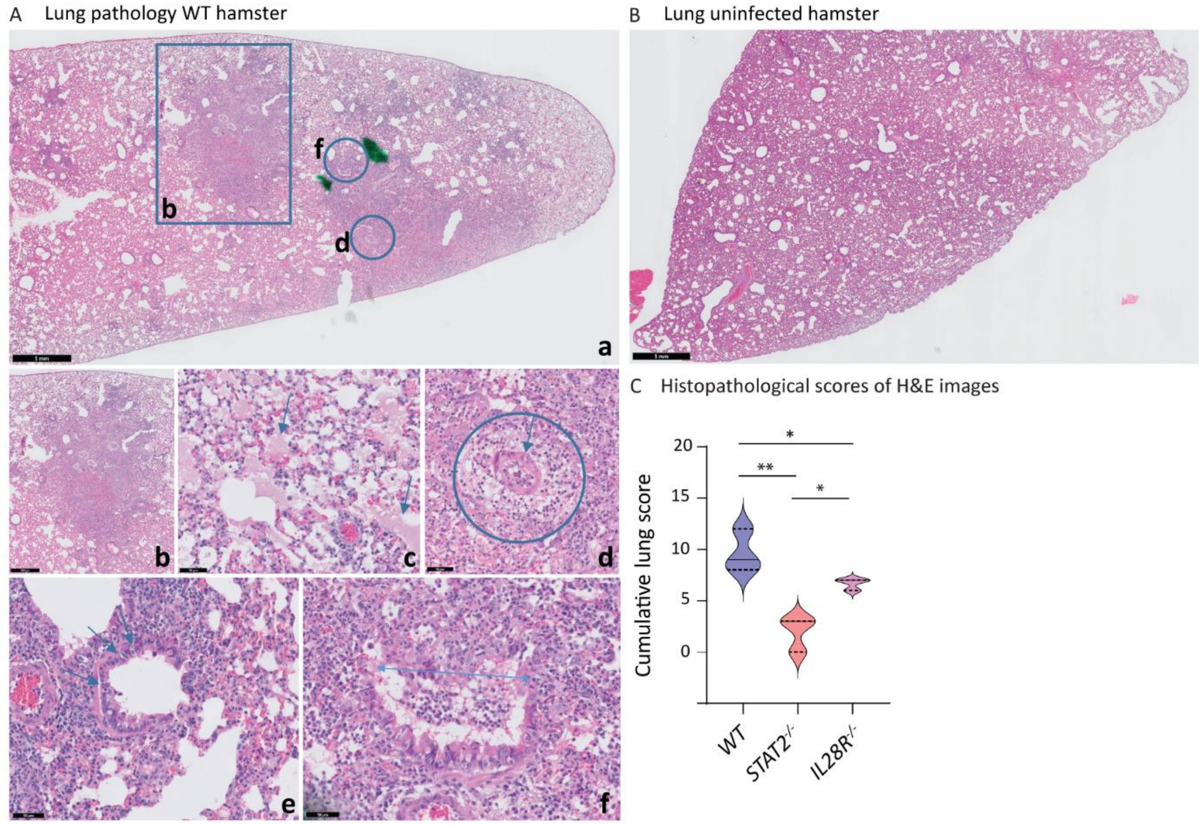


**Suppl Fig. S8. Lung pathology in a WT hamster**. **(A)** Representative H&E images of the lung of a P4 SARS-CoV-2 infected WT hamster. **(Panel a)** rectangle bronchopneumonia (see also panel b), upper circle necrotizing bronchiolitis (see also panel f), lower circle perivascular edema (see also panel d). **(Panel b)** area of bronchopneumonia with inflammation centered around a bronchus (magnification of panel a). **(Panel c)** intra-alveolar hemorrhage and edema (arrows). **(Panel d)** perivascular edema (circle), vascular wall (arrow) is surrounded by loose fibrous tissue and a few inflammatory cells (magnification of panel a). **(Panel e)** bronchus with apoptotic cells (arrows). **(Panel f)** necrotizing bronchiolitis: half of the bronchus wall has disappeared and is replaced by inflammatory cells (arrow, magnification of panel a). **(B)** Representative H&E image of the lung of an uninfected WT hamster. **(C)** Violin plot of the cumulative clinical scores of lung damage in WT (n = 4), *STAT2*$^{-/-}$ (n = 4) and *IL28R-a*$^{-/-}$ (n = 4) hamsters (apoptotic bodies, necrotizing bronchiolitis, edema, pneumonia and inflammation) are indicated. Statistical significance between groups was calculated by the nonparametric two-tailed Mann Whitney U-test (* P < 0.05, ** P < 0.01).

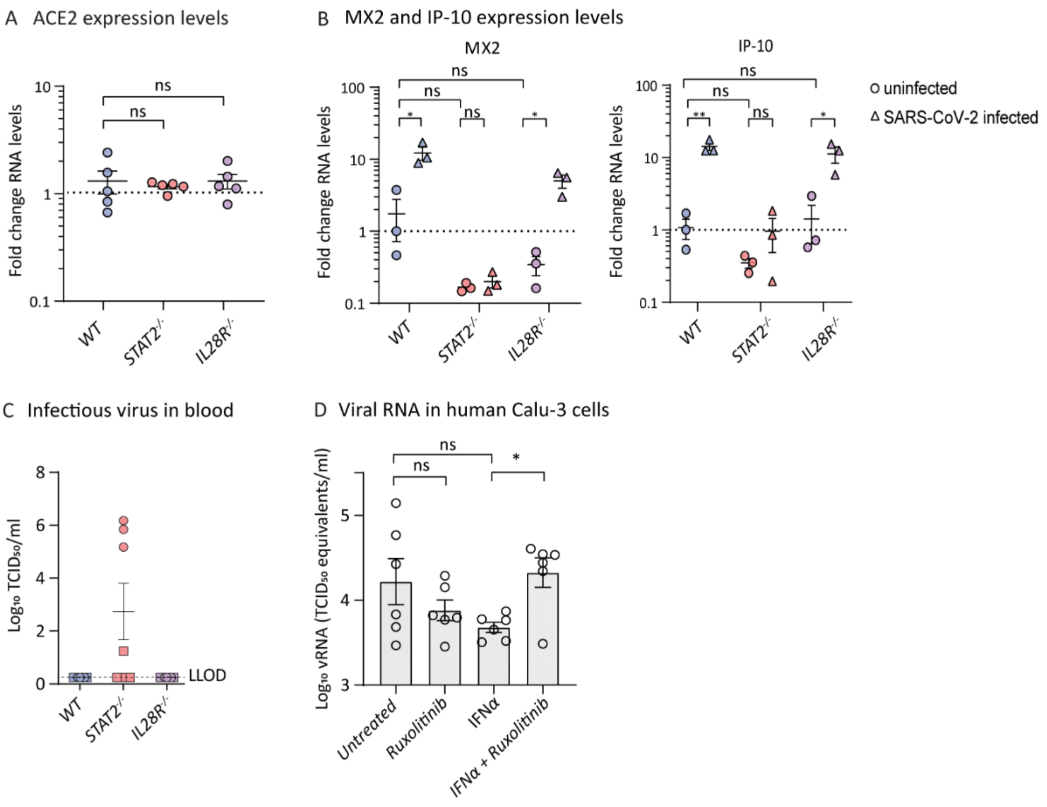


**Suppl Fig. S9. Validation of the hamster model. (A)** *ACE2* RNA expression levels in lungs of WT (n=5), *STAT2*$^{-/-}$ (n=5) and *IL28R-a*$^{-/-}$ (n=5) hamsters. **(B)** *MX2* (induced by IFNα/STAT2 signaling) and *IP-10* (induced by both type I and type II IFNs) RNA expression levels in lungs of uninfected (circles) or SARS-CoV-2-infected (triangles) (day 4 p.i.) WT (2× n=3), *STAT2*$^{-/-}$ (2× n=3) and *IL28R-a*$^{-/-}$ (2× n=3) hamsters. **(C)** Infectious viral loads in the blood of SARS-CoV-2-infected WT, *STAT2*$^{-/-}$ and *IL28R-a*$^{-/-}$ hamsters. Outcomes derived from inoculation with passage 4 or passage 6 SARS-CoV-2 is designated by circles (P4) or squares (P6). Infectious viral loads in the blood are expressed as the number of infectious virus particles per ml of blood. **(D)** Enhancement of SARS-CoV-2 replication in human airway epithelial cells by pharmacological ablation of STAT signaling. Calu-3 (human airway epithelial) cells[10] were left untreated or treated with Ruxolitinib (4 µM), Type I IFN (10 IU/mL) or a combination of both. Treatment was initiated 4 h before infection and was continued through the whole experiment[7]. Cultures were infected with SARS-CoV-2 at a MOI of 1. At 48 h p.i. cell culture SN was collected, RNA was extracted and the amount of vRNA was quantified using RT-qPCR. A serial dilution of the same virus stock was prepared in medium and processed in the same manner to generate a standard curve for absolute quantification. Inhibiton of JAK/STAT signaling by Ruxolitinib[11] can rescue SARS-CoV-2 virus replication from the antiviral effect of Type-I IFN. One-way ANOVA with post hoc test for multiple comparisons was performed to compare groups as indicated. P-values in are shown as * ≤ 0.05.

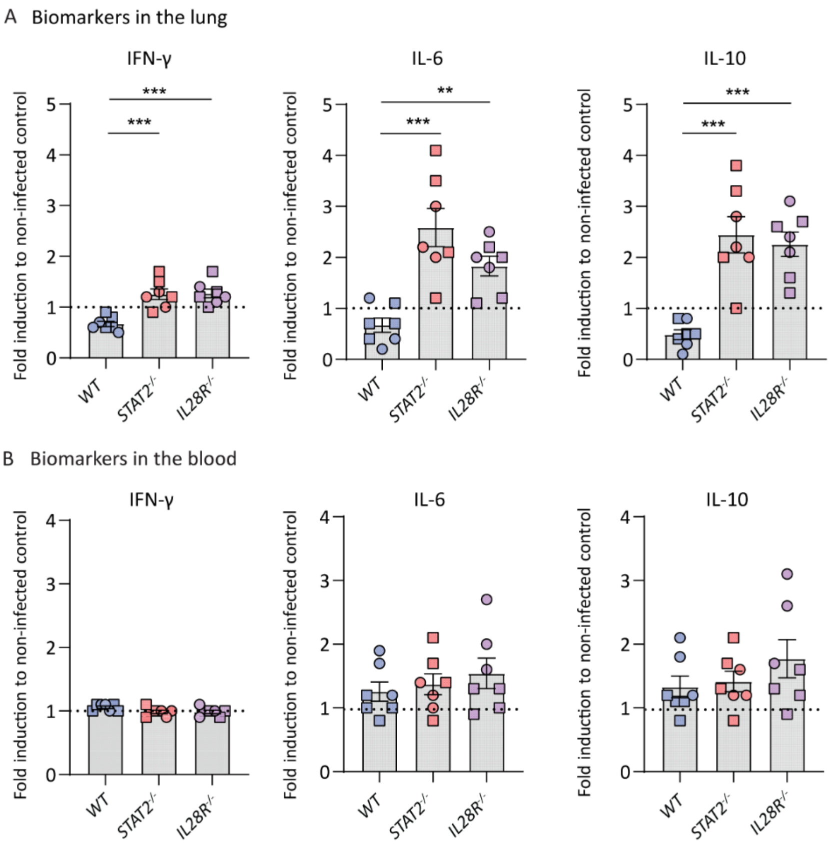


**Suppl Fig. S10. Tissue- and blood-based biomarkers of lung infection and inflammation in SARS-CoV-2 infected hamsters**. Outcomes derived from inoculation with passage 4 or passage 6 SARS-CoV-2 is designated by circles (P4) or squares (P6). Levels of IFN-γ, IL-6 and IL-10 in **(A)** lung homogenates and **(B)** serum of SARS-CoV-2 infected WT, *STAT2*$^{-/-}$ and *IL28R-a*$^{-/-}$ hamsters relative to non-infected controls of the same strain as determined by ELISA. Values for infected animals (n=7 each) compiled from two independent experiments using either P4 (n=3) and P6 (n=4) virus, relative to non-infected controls of the same strain (n=3 each). Statistical significance between groups was calculated by the nonparametric two-tailed Mann-Whitney U-test (** $P < 0.01$, *** $P < 0.001$).

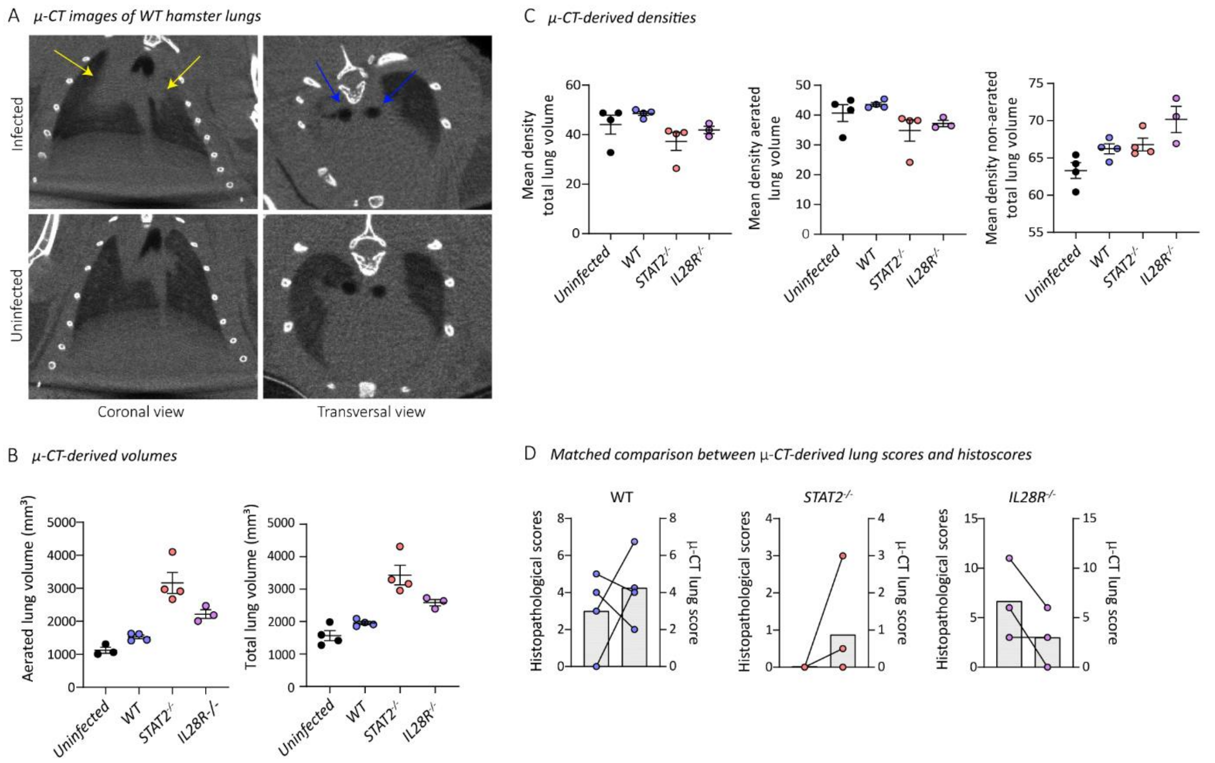


**Suppl Fig. S11. Imaging-based biomarkers of micro-CT-observed lung pathology. (A)** Representative coronal and transversal micro-CT (μCT) images of an upper airway enlargement and dense lung infiltrates in the lungs of infected (P6 SARS-CoV-2) WT hamsters compared to uninfected hamsters. Arrows indicate examples of pulmonary infiltrates seen as consolidation of lung parenchyma (yellow), or upper airways dilatations (blue). Micro-CT-derived biomarkers quantifying aerated lung volume and total lung volume **(B)** and corresponding mean densities therein **(C)** (see also Fig. 3D). The overall increase in aerated lung volume (healthy lung tissue) and associated increase in total lung volume for the *STAT2*$^{-/-}$ hamsters is likely due to a higher age of the hamsters used in the study (seven- to twelve-week old *STAT2*$^{-/-}$ hamsters versus five- to seven-week-old *IL28R-a*$^{-/-}$ hamsters). **(D)** Matched comparison between μCT-derived semi-quantitative lung scores (left Y-axis) **(values from Fig. 3B)** and histoscores (right Y-axis) (values from Fig. 2D). Lines indicate matched samples. The data shown are means ± SEM.

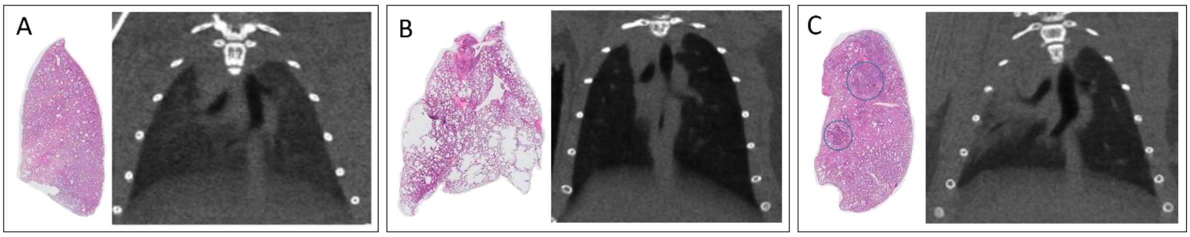


**Suppl Fig. 12. Correspondence between *in vivo* micro-CT and *ex vivo* histopathological observations.** **(A)** Histopathological H&E-stained section of whole lobe (left) and coronal micro-CT image of a healthy, uninfected hamster lung. **(B)** Histopathology confirmed hyperinflation visualized on the micro-CT scan as decreased density (blackening) of lung tissue and overall lung enlargement, seen in one infected *STAT2*$^{-/-}$ hamster. **(C)** Consolidation of right lung parenchyma on micro-CT corresponds to bronchopneumonia evidenced by histopathology (darker blue stained areas, blue circles) of an infected IFNL-KO hamster. Micro-CT also shows dilation of the upper airways.

**Extra references to Supplementary Figures**

**Supplementary Table S1**. Primers and probes used for RT-qPCR

| Gene | Description | Oligonucleotide sequence |
|---|---|---|
| **SARS-CoV-2** | Forward primer | 5'-TTA CAA ACA TTG GCC GCA AA-3' |
| | Reverse primer | 5'-GCG CGA CAT TCC GAA GAA-3' |
| | Probe | 5'-FAM-ACA ATT TGC CCC CAG CGC TTC AG-BHQ1-3' |
| **Mouse *Actb*** | Forward primer | 5'-GAT TAC TGC TCT GGC TCC TAG-3' |
| | Reverse primer | 5'-GAC TCA TCG TAC TCC TGC TTG-3' |
| | Probe | 5'-FAM-CTG GCC TCA CTG TCC ACC TTC C-ZEN/IABkFQ-3' |
| **Hamster *MX2*** | Forward primer | 5'-CCA GTA ATG TGG ACA TTG CC-3' |
| | Reverse primer | 5'-CAT CAA CGA CCT TGT CTT CAG TA-3' |
| | Probe | 5'-FAM-TGT CCA CCA GAT CAG GCT TGG TCA-ZEN/IABkFQ-3' |
| **Hamster *IP-10*** | Forward primer | 5'-GCC ATT CAT CCA CAG TTG ACA-3' |
| | Reverse primer | 5'-CAT GGT GCT GAC AGT GGA GTC T-3' |
| | Probe | 5'-FAM-CGT CCC GAG CCA GCC AAC GA-ZEN/IABkFQ-3' |
| **Hamster *ACE2*** | Forward primer | 5'-GGG AAC TGT CAA AGG GTA CAG-3' |
| | Reverse primer | 5'-CCC TTC CTA CAT CAG TCC TAC T-3' |
| | Probe | 5'-TCC CTG CTC ATT TGC TTG GTG ACA-ZEN/IABkFQ-3' |
| **Hamster *ACTB*** | Forward primer | 5'-GGC CAG GTC ATC ACC ATT-3' |
| | Reverse primer | 5'-GAG TTG AAT GTA GTT TCG TGG ATG-3' |
| | Probe | 5'-Cy5-TTT CCA GCC TTC CTT CCT GGG TAT G-IBRQ-3' |

**Supplementary Table S2.** RT-qPCR Taqman assays for analysis of DEG in SARS-CoV-2 infected cells

| **Gene of Interest** | **Proposed MoA in immune response** | **Reference** | **Assay ID** |
|---|---|---|---|
| *18s rRNA* | House keeping gene | | |
| *GAPDH* | House keeping gene | | |
| *AIM2* | Cytosolic DNA sensor | Yu and Levine[1] | Mm01295719_m1 |
| *AKT1* | Controls innate immune cell development and function | Zhang et al[2] | Mm01331626_m1 |
| *CHUK (IKK)* | Inhibitor of nuclear factor kappa-B kinase subunit alpha (IKK-α) | Llona-Minguez et al[3] | Mm00432529_m1 |
| *CXCL11* | Interferon inducible T cell alpha chemoattractant | Venter et al[4] | Mm00444662_m1 |
| *DHX36* | Enhances RIG-I Signaling | Yoo et al[5] | Mm00661002_m1 |
| *DDX58* | Antiviral, IFN signalling, innate immune receptor | Pulendran et al[6], Querec et al[7] | Mm01216853_m1 |
| *EIF2AK 2* | Antiviral, IFN signalling | Querec et al[7] Gaucher et al[8] | Mm01235643_m1 |
| *IFI16* | Interferon Gamma Inducible Protein 16 | Trapani et al[9] | Mm00492602_m1 |
| *IFIH1* | Antiviral, IFN signalling, innate immune receptor | Pulendran et al[6] Querec et al[7] | Mm00459183_m1 |
| *IFT122* | Intraflagellar Transport 122 | Boubakri et al[10] | Mm00661643_m1 |
| *IRF3* | Interferon regulatory factor 3 | Collins et al[11] | Mm00516784_m1 |
| *IRF9* | Antiviral, IFN signalling | Gaucher et al[8] | Mm00492679_m1 |
| *MB21D1(cGAS)* | Cytosolic DNA sensor | Liang et al[12] | Mm01147496_m1 |
| *MRE11* | Double strand break repair nuclease | Shibata et al[13] | Mm00450600_m1 |
| *OAS1* | Antiviral, IFN signalling | Gaucher et al[8] | Mm00449297_m1 |
| *OAS3* | Antiviral, IFN signalling | Gaucher et al[8] | Mm00460944_m1 |
| *MX1* | Antiviral, IFN signalling | Querec et al[7], Gaucher et al[8] | Mm00487796_m1 |
| *PDK1* | Key role in regulation of glucose and fatty acid metabolism | Tan et al[14] | Mm00554300_m1 |
| *PDPK1* | Important role in the signalling pathways | Sato et al[15] | Mm00440707_m1 |
| *PLSCR1* | Antiviral, IFN signalling | Querec et al[7], Gaucher et al[8] | Mm01228223_g1 |
| *PNPT1* | Antiviral, IFN signalling , innate immune receptor | Pulendran et al[6], Querec et al[7] | Mm00466286_m1 |

| | | | |
|---|---|---|---|
| *STAT1* | Key regulators of the early innate immune response | Decker et al[16] | Mm01257286_m1 |
| *STAT3* | Control of inflammation and immunity | Hillmer et al[17] | Mm01219775_m1 |
| *TBK1* | Protein kinase downstream of TMEM173 (STING) | Lam et al[18] | Mm00451150_m1 |
| *TLR3* | pattern-recognition receptors (PRRs) | Yu and Levine[1] | Mm01207404_m1 |
| *TLR9* | Cytosolic DNA sensor | Yu and Levine[1] | Mm07299609_m1 |
| *TMEM173(STING)* | Stimulator of IFN response | Lam et al[18] | Mm01158117_m1 |
| *TP53* | Tumor protein 53 | Matlashewski et al[19] | Mm00480750_m1 |
| *TRAF6* | TNF receptor associated factor (TRAF) | Youseff et al[20] | Mm00493836_m1 |
